# SifA-mediated Remodeling of the *Salmonella*-Containing Vacuole Prevents Bacterial Dormancy by Promoting Nutrient Accessibility

**DOI:** 10.1101/2025.01.08.631959

**Authors:** Camila Valenzuela, Francisco J. Garcia Rodriguez, Thomas Mazza, Magdalena Gil, Kateryna Dotsenko, Chak Hon Luk, Jost Enninga

## Abstract

*Salmonella* exploits a range of intracellular niches within host cells, adapting to different microenvironments that support its survival and replication. This niche diversity is mediated by bacterial effectors injected via the two type 3 secretion systems (T3SS). *Salmonella* resides either within membrane-bound *Salmonella* containing vacuoles (SCV) or they grow rapidly within the cytosol upon SCV rupture. Recently, we identified a third intracellular subpopulation, dormant *Salmonella* within modified vacuoles in epithelial cells, which can survive for up to 7 days *in vitro*. To explore how bacterial effectors influence the balance of these three subpopulations, we constructed a panel of mutants lacking genes encoding key effectors and examined their intracellular behaviors at 6 and 24 hours post-infection (hpi). Deletion of the T3SS-2 effector SifA significantly increased the dormant subpopulation at later infection time points, identifying SifA as the first known effector regulating bacterial dormancy in epithelial cells. SifA-induced *Salmonella*-induced filaments (SIFs) characterize the mature *Salmonella*-containing vacuole (SCV), and we observed that SIFs were needed for intravacuolar growth of *Salmonella*. We could show that SIF formation was critical for nutrient acquisition within the SCV; inhibition of SIF formation resulted in a higher proportion of dormant bacteria, akin to the effect of reduced glucose availability during infection. Both bacterial and host glycolysis pathways were required to prevent dormancy, as proper nutrient scavenging through SIF-mediated modification of the SCV is essential for maintaining a replicative, non-dormant state. These results underscore the importance of nutrient access, facilitated by reprogramming host endomembrane trafficking, for *Salmonella* to avoid dormancy and sustain active replication within its intracellular niche.

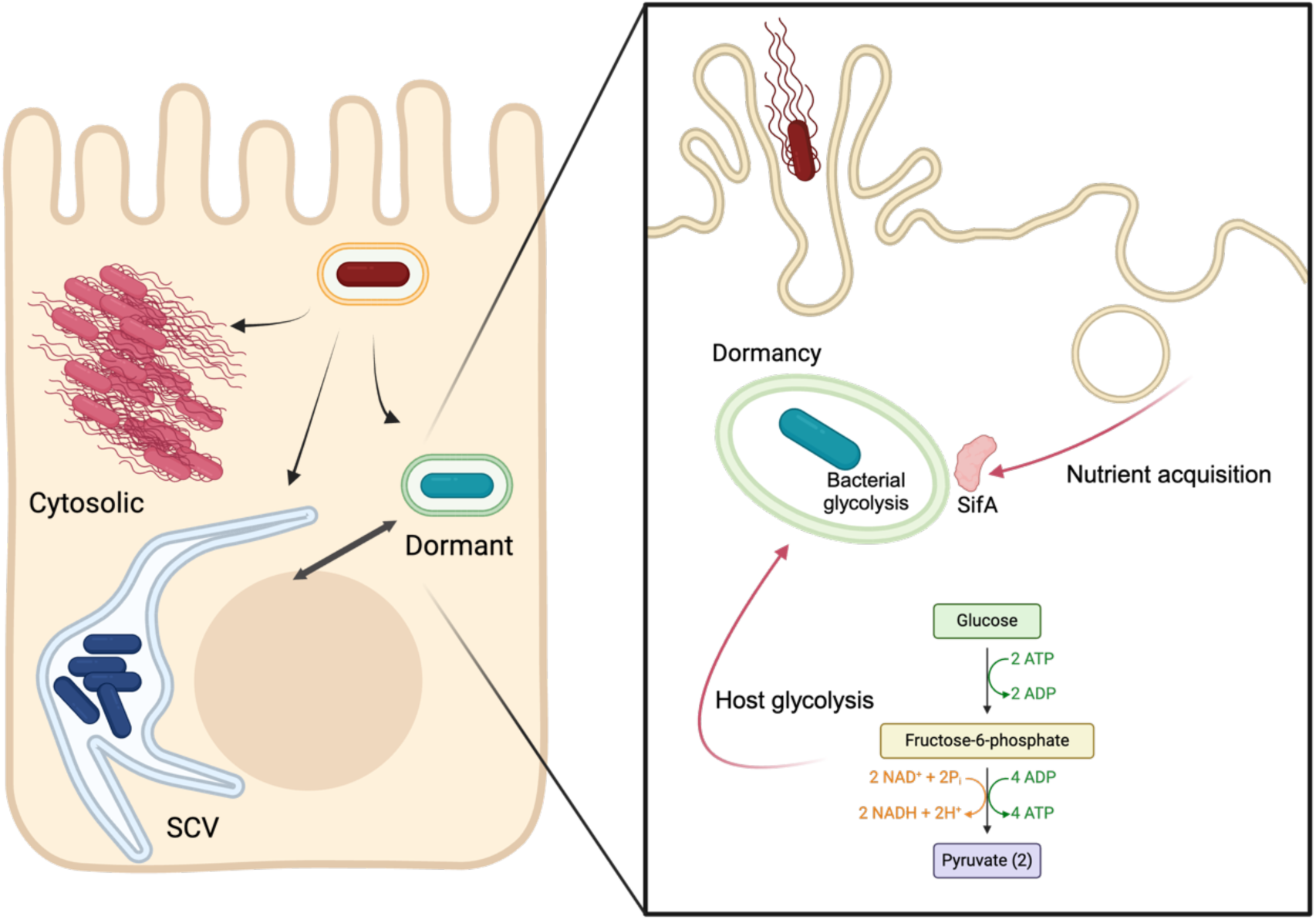

## Introduction

*Salmonella enterica* serovar Typhimurium is a major pathogen that causes gastrointestinal infections and systemic disease in humans and animals^1^. Upon ingestion, the bacterium interacts with epithelial cells of the small intestine entering them and surviving within them^2^. *Salmonella* induces its internalization into non-phagocytic epithelial cells from the apical side, using a type 3 secretion system (T3SS) encoded in the *Salmonella* Pathogenicity Island 1 (SPI-1), T3SS-1, that injects a cocktail of bacterial effectors to take control of several host pathways^3^. Once internalized, *Salmonella* is contained inside a vacuolar compartment named *Salmonella*-containing vacuole (SCV). This SCV is modified by effectors injected by a second T3SS-2, encoded in SPI-2, that allows the generation of a replicative niche. The maturation of this compartment comprises the acquisition of late endosome markers, such as RAB7 and LAMP-1. Remarkably, *Salmonella* can colonize other intracellular niches in epithelial cells. Upon rupture of the SCV, the bacterium is either recognized by the host cell and targeted by the autophagy machinery leading to its degradation, or it escapes this recognition and starts replicating in the host cytosol at a very high rate, in what has been named hyper-replicating cytosolic *Salmonella*^4,5^. In addition to these already somewhat characterized intracellular lifestyles, a novel subpopulation inside epithelial cells in a dormant state has recently been described by us^6^. Using correlative light and electron microscopy (CLEM) we were able to determine that dormant *Salmonella* resides within a vacuolar compartment, but importantly, this vacuole does not show the classical vacuolar markers of a canonical SCV, with only minor LAMP1 recruitment^6^. Importantly, dormant *Salmonella* within these modified vacuoles are more tolerant to antibiotic treatment, show very low metabolic activity and can persist for up to 7 days inside cells *in vitro*.

One of the hallmarks of SCV maturation is the generation of networks of *Salmonella*- induced tubules (SITs) that extend outward from the SCV and spread throughout the host cell^7^. Among these, the *Salmonella*-induced filaments (SIFs) are the most prevalent and well-characterized type of SIT^8,9^. These structures have the same membrane composition as the mature SCV enriched in the late endosome markers LAMPs, RAB7, cholesterol and the vATPase, while lacking the characteristic late endosomal/lysosomal markers cathepsin D, lysobisphosphatidic acid (LBPA) and mannose-6-phosphate receptor (M6PR)^10–12^. Multiple studies have demonstrated fusion events between compartments derived from the endocytic pathway and the SIF network, leading to the acquisition of the mentioned membrane markers^9,11,13^. Moreover, there is evidence of exchange of luminal contents between the SCV and the SIF network, that would explain the access to endocytosed material^7,9^. Significantly, *Salmonella* in vacuoles with SIFs are more metabolically active than those without SIFs, suggesting a role for SIFs in nutrient acquisition^9^.

SIF formation depends on the activity of T3SS-2 effectors, in particular PipB2, SifA, SopD2, SseF, SseG and SseJ^14^. Among them, SifA has been recognized as the key regulator^15^, as demonstrated by the inability of *sifA* mutants to generate SIFs in HeLa cells^15^. Besides their inability to form SIFs, *sifA* mutants were associated with vacuolar rupture and originally described as preferentially cytosolic. Nevertheless, the efficiency of cytosolic access has remained unclear, and it can only be observed several hours upon host cell challenge. SifA-mediated cytosolic access has been explained by the instability of the vacuole when SifA is absent, due to the excessive recruitment of kinesin-1, and the activity of other effectors such as SseJ and SopD2^16^.

In different types of epithelial cell lines, around 10% of the infected cells contain cytosolic *Salmonella*^4^. Once in the cytoplasm, *Salmonella* growth occurs at a very high rate, reaching hundreds of bacteria per cell, in what has been termed hyper-replicating *Salmonella*^5^. Importantly, cytosolic *Salmonella* expresses the T3SS-1 and flagella, rendering them primed for infection if they reach neighboring cells^5^.

*Salmonella* can enter a dormant state within various host cell types, and the establishment of intracellular niches of targeted host cells is a critical aspect of the pathogen’s survival and virulence. *Salmonella* can form non-replicating persisters in macrophages, a dormant state that contributes to chronic infections^17–19^. These dormant bacteria remain viable and may express virulence factors at later stages, and dormancy-related factors have been studied in more detail almost exclusively in this cell type. We recently discovered a subpopulation of dormant *Salmonella* in epithelial cells, these bacteria reside in vesicular compartments distinct from conventional SCVs^6^. The specific mechanisms and niches associated with dormancy in epithelial cells are not well characterized and may involve several bacterial and host pathways.

## Results

### *Salmonella* effectors mutants exhibit altered intracellular niche distributions

Given the diversity of intracellular niches, we leveraged our previously published SINA reporters^6^ to investigate factors contributing to intracellular niche formation. We first evaluated the contribution of specific *Salmonella* effectors to the subpopulation distribution within infected epithelial cells at 6 and 24 hours post-infection (hpi), differentiating between vacuolar, cytosolic and dormant niches. For this, we constructed a collection of deletion mutants (**Table S1**) targeting the main effectors required for *Salmonella* to modify the SCV and screened them using our SINA readout in HeLa cells (**Figure 1B-E** and **Figure S1**). We observed that deleting single effectors led already to noticeable effects on the level of cells containing cytosolic *Salmonella* at 6 hpi, with τι*sifA* and τι*sopF* mutants showing an increase in this subpopulation (**Figure 1B-C** and **Figure S1**). These findings echoed previous reports portraying these two effectors as gatekeepers of the host cytosol^12,16,20^. Despite this, we noted that only a minority of cells invaded by these mutants showed cytosolic bacteria suggesting that the gatekeeper function does not fully characterize the action of these effectors. Furthermore, no significant effects were observed for the dormant subpopulation at this early time point (**Figure 1D**). In contrast, the dormant subpopulation was highly affected at the later 24 hpi timepoint, with several mutants altering the level of cells containing dormant *Salmonella* while diminishing the levels of vacuolar *Salmonella* (**Figure 1C-1E** and **Figure S1**). One of the most striking dormancy phenotypes was observed in cells invaded with the τι*sifA* mutant at 24 hpi, where up to 40% of the infected cells harbor dormant *Salmonella*.

**Figure 1.**
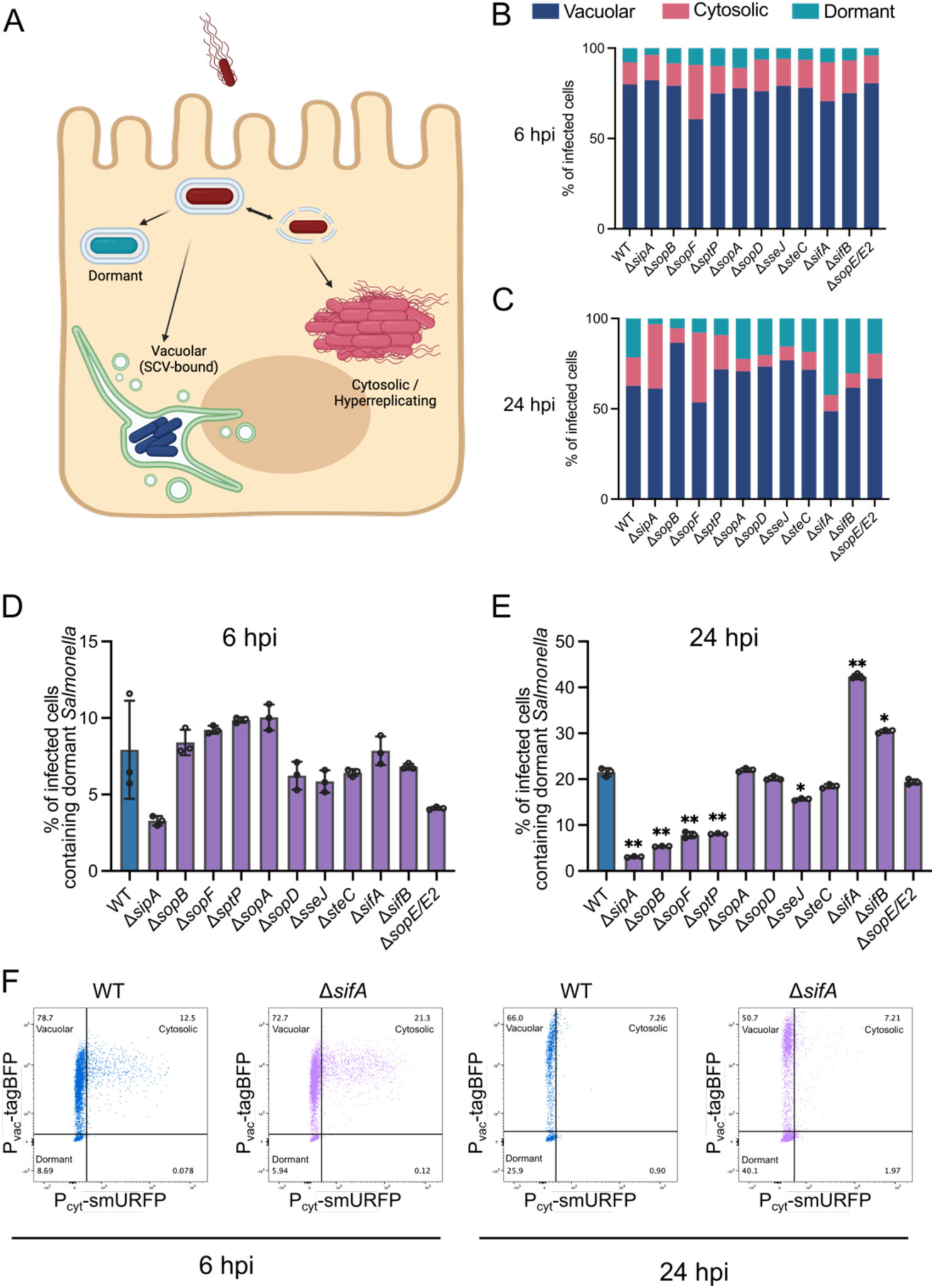
*Salmonella* effector mutants show altered intracellular niche formation: **(A)** Graphical illustration of the different intracellular niches colonized by *Salmonella* expressing SINA1.1 in HeLa epithelial cells. Intracellular subpopulation distribution at 6 **(B)** and 24 hpi **(C)**, the percentage of infected cells containing each subpopulation is represented in blue (vacuolar), pink (cytosolic) and dormant (turquoise), for all strains. The percentage of cells containing dormant *Salmonella* at 6 **(D)** and 24 hpi **(E)** of the different deletion mutants (purple) compared with the WT strain (blue) is shown. **(F)** Representative flow cytometry plots of HeLa cells infected with *S*. Typhimurium WT or τι*sifA* at 6 and 24 hpi. Bars show the average of 3 biological replicates (N=3). Statistical significance was evaluated using a one-way ANOVA followed by a with Dunnett’s multiple comparison test, * = *P*<0.05, ** = *P*<0.01.

### SIF formation correlates with lower dormancy in epithelial cells

The effector SifA is involved in one of the main structural modifications of the SCV that is most relevant for intravacuolar nutrition: the generation of the SIF network^9^. Therefore, we first evaluated if the dormant vacuole shows the presence of SIFs at 6 hpi, and observed that contrary to vacuolar *Salmonella* showing typical growth for this compartment, the dormant ones reside in compartments devoid of SIFs (**Figure 2A-B**), while still positive for RAB7 (**Figure S2**). We previously reported that these dormant vacuoles are negative for LC3 and therefore not targeted by the autophagy machinery^6^. Additionally, we used an image-based analysis to evaluate intracellular niche formation over time, along with the presence of SIFs. For this, we used HeLa cells and *Salmonella* expressing SINA1.1 or HeLa LAMP1-GFP cells and *Salmonella* expressing SINA1.7 and added nocodazole, an inhibitor of microtubule polymerization, at 1 hpi to inhibit SIF growth. To track and quantify SIF formation in HeLa LAMP1-GFP cells we developed an image-analysis pipeline (**Figure S3**), in the presence and absence of nocodazole or different strains. We were able to confirm that only the WT strain generated SIFs, and the formation of them is inhibited using nocodazole (**Figure 2C**). By tracking the appearance of dormant bacteria in infected HeLa cells in the presence or absence of nocodazole, we were able to determine that inhibiting SIF formation using nocodazole, increased the number of dormant bacteria (**Figure 2D-E**). Moreover, the τι*sifA* mutant exhibited faster kinetics in terms of the appearance of dormant *Salmonella* compared with the WT strain (**Figure 2D**). Using our analysis pipeline, we were able to quantify at the single-bacterium level the appearance of each bacterial subpopulation (**Figure S4A**), showing that the different intracellular niches have different patterns of growth. Moreover, by analyzing the Timer^510/580^ ratio as a readout of the replication rate of each individual bacterium we observed that at the single-bacterium level the presence of nocodazole lowers the growth efficiency in the vacuolar and cytosolic subpopulations (**Figure S4B**). Together, our results indicate that the SIF network helps prevent *Salmonella* from going into dormancy.

**Figure 2.**
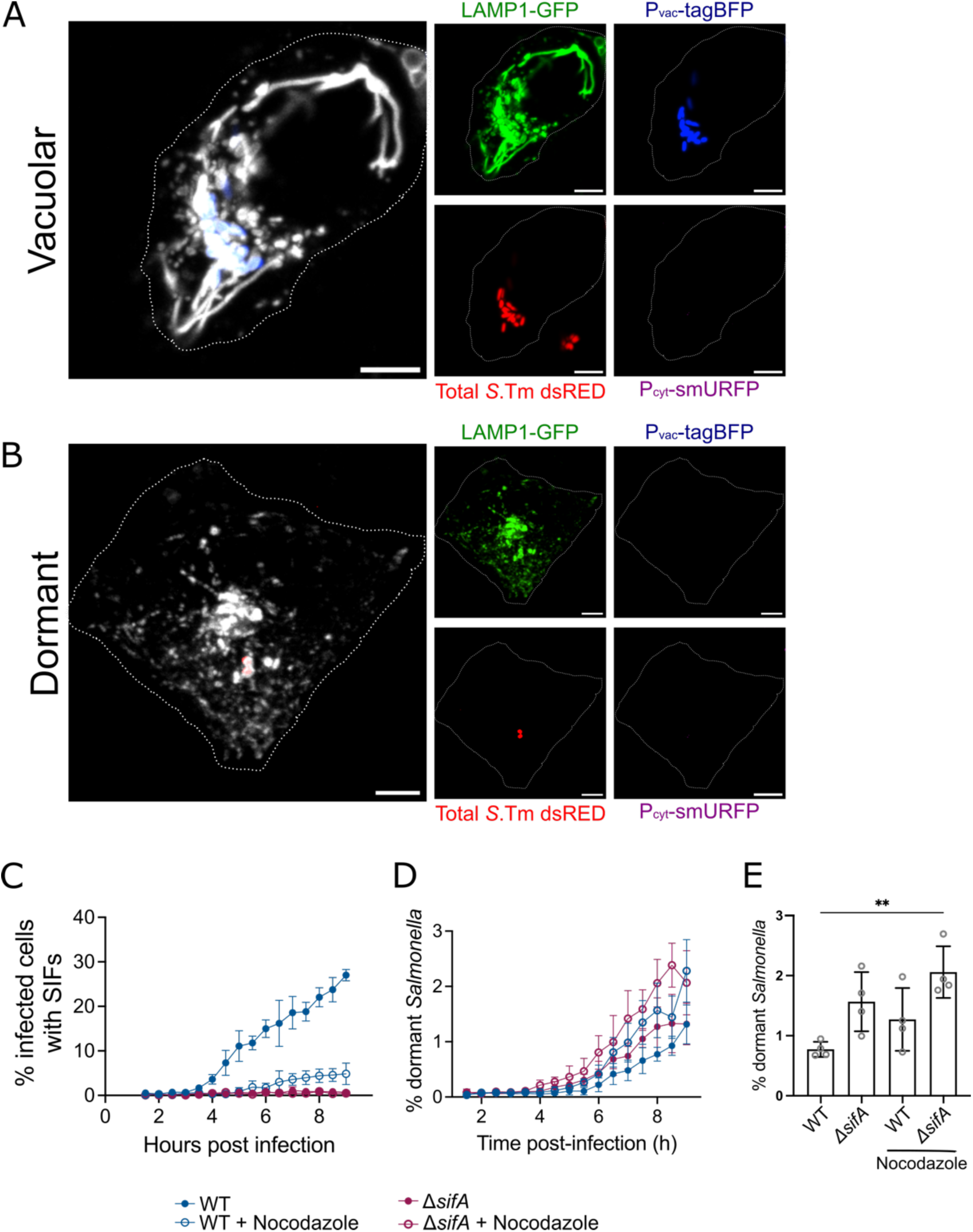
Inhibition of SIF formation is correlated with *Salmonella* dormancy in epithelial cells. Representative confocal images of LAMP1-GFP-expressing HeLa cells infected with *Salmonella* expressing SINA1.7 growing in a SCV **(A)** or in a dormant state **(B)**. Green: LAMP1-GFP, red: dsRED total *Salmonella*, blue: vacuolar marker Pvac-tagBFP, magenta: cytosolic marker P_cyt_-smURFP. Merge shows the LAMP1-GFP distribution and *Salmonella* according to its intracellular niche (blue: vacuolar, red: dormant). **(C)** Quantification using a custom pipeline of infected cells with SIFs, separated by subpopulation over time (see methods and **Figure S3**), in untreated or nocodazole 1 1M treated cells. **(D)** Quantification of dormant *Salmonella* in treated and untreated cells over time and **(E)** at 6 hpi.

### Nutrient accessibility and the glycolytic state of the cell influence the generation of dormant *Salmonella*

To further evaluate the role of intravacuolar nutrition of the invading pathogen, we performed infections using different glucose concentrations in the cell culture medium during infection. Glucose is the main carbon source used by *Enterobacteria* such as *S*. Typhimurium and glycolysis is necessary to allow efficient intracellular growth in the SCV^21^. We determined when using low glucose media, *Salmonella* had a greater tendency to become dormant in HeLa cells at 6 hpi, for both the WT and τι*sifA* strains (**Figure 3**). At 24 hpi, low glucose conditions increased the level of cells containing dormant bacteria (**Figure 3B-3D**). It has been reported that cell culture media nutrients are available to intravacuolar *Salmonella* in RAW264.7 macrophages in a T3SS-2- dependent manner^9^.

**Figure 3.**
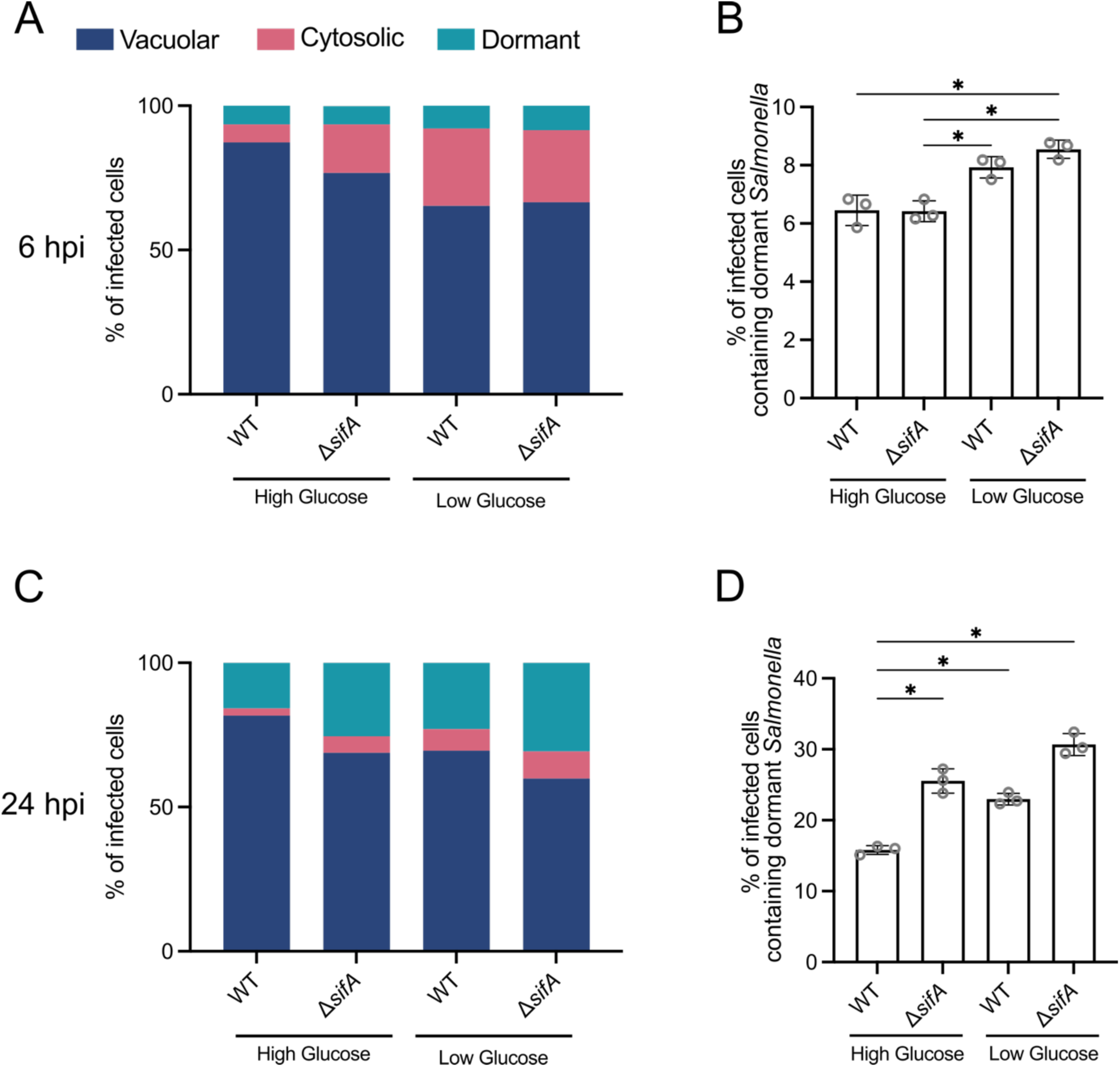
Lower nutrient availability correlates with an increase of *Salmonella* dormancy in epithelial cells. Intracellular subpopulation distribution at 6 **(A)** and 24 hpi **(C)**, the percentage of infected cells containing each subpopulation is represented in blue (vacuolar), pink (cytosolic) and dormant (turquoise). The percentage of cells containing dormant *Salmonella* at 6 **(B)** and 24 hpi **(D)** in high (25 mM) or low (2.8 mM) glucose concentration is shown. Plots show the average of 3 biological replicates (N=3). Statistical significance was evaluated using a one-way ANOVA followed by a Tukey’s multiple comparison test, * = *P*<0.05.

These results indicate that the accessibility of nutrients in the extracellular environment is relevant for intravacuolar *Salmonella* with glycolysis being a relevant host pathway in epithelial cells. To further explore this possibility, we first performed infections in cells pre-incubated in media containing galactose instead of glucose, as utilization of this sugar as carbon source inhibits glycolysis in epithelial cells^22^. Our results indicate that in cells growing in galactose, the number of cells containing dormant *Salmonella* increased dramatically at 6 hpi for both the WT and the τι*sifA* strains (**Figure 4A-B**). We confirmed these results using 2-Deoxy-D-glucose (2DG), a glycolysis inhibitor that competitively inhibits the phosphoglucoisomerase, therefore preventing the formation of glucose-6-phosphate. Importantly, to decouple host and bacterial glycolysis, we performed the experiments in a τι*manZ* background, as this mutant is unable to import 2DG, rendering *Salmonella* insensitive to this type of glycolysis inhibition^23^. Using the 2DG inhibitor, we observed a significant increase in cells containing dormant *Salmonella* at 6 hpi (**Figure 4C-D**), independently of SifA, in accordance with our initial screening (**Figure 1**). At 24 hpi, the addition of 2DG increased the levels of cells containing dormant *Salmonella* in the WT (τι*manZ*) compared with control cells (**Figure 4E-F**). Deleting *sifA* had no additional effect in dormancy formation compared with untreated cells (**Figure 4F**). Altogether, our data indicates that the glycolytic state of the cell is relevant in the host-pathogen dynamics, as inhibition of host glycolysis increases the levels of dormant *Salmonella* in epithelial cells.

**Figure 4.**
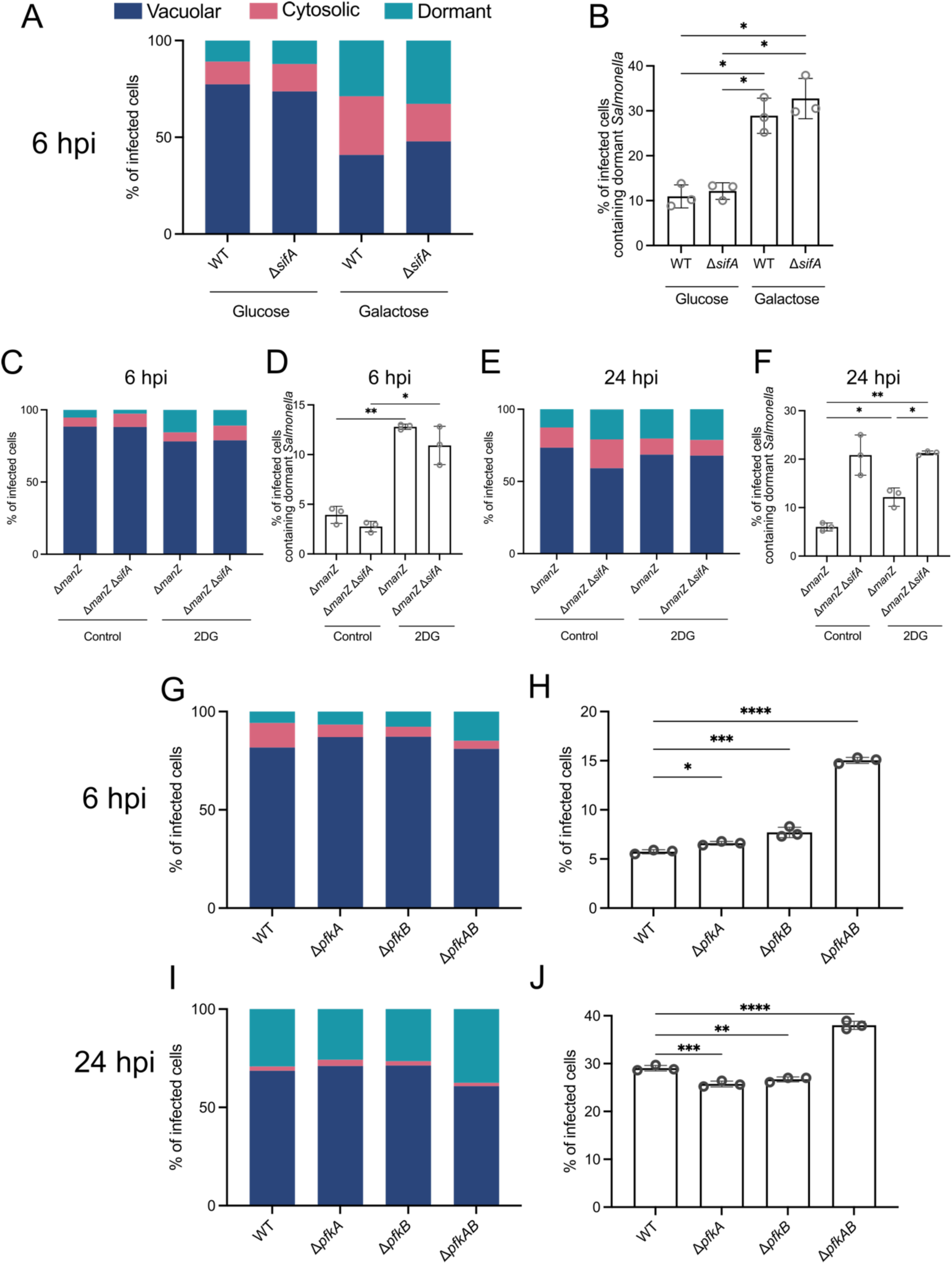
Host and bacterial glycolysis are involved in *Salmonella* dormancy in epithelial cells. The percentage of infected cells containing each subpopulation is represented in blue (vacuolar), pink (cytosolic) and dormant (turquoise). **(A)** Inhibition of host-cell glycolysis by modifying the cell media carbon source from glucose 25 mM to galactose 10 mM changes the intracellular niche distribution. **(B)** The use of galactose increases the percentage of cells containing dormant *Salmonella* at 6 hpi. **(C-F)** Inhibition of host-cell glycolysis using 2-deoxy-D-glucose (2DG) in a τι*manZ* background, alters intracellular niche distribution **(C** and **E)** and increases the percentage of cells containing dormant *Salmonella* **(D** and **F)** at 6 and 24 hpi. Statistical significance was evaluated using a one-way ANOVA followed by a Tukey’s multiple comparison test, * = *P*<0.05, ** = *P*<0.01. **(G-H)** Abolishing bacterial glycolysis by deleting the genes encoding the two isoforms of the phosphofructokinase (*pfk*) increases the formation of dormant *Salmonella* in epithelial cells. Intracellular subpopulation distribution at 6 **(G)** and 24 hpi **(I)** of the single and double mutants compared with the WT strain. Percentage of cells containing dormant *Salmonella* at 6 **(H)** and 24 hpi **(J)**. Statistical significance was evaluated using a one-way ANOVA followed by a with Dunnett’s multiple comparison test * = *P*<0.05, ** = *P*<0.01*** = *P*<0.001, **** = *P*<0.0001. Plots show the average of 3 biological replicates (N=3).

### *Salmonella* requires a functional glycolysis pathway to avoid dormancy

To evaluate the contribution of bacterial glycolysis to dormancy, we deleted the genes encoding the two phosphofructokinase isoforms in *Salmonella*, τι*pfkA* and τι*pfkB*^21,24^. We then performed infections in a high glucose condition and evaluated the level of cells containing dormant bacteria at 6 and 24 hpi. Our results show that the τι*pfkAB* double mutant presented a significant increase in the level of intracellular dormant bacteria (**Figure 4G-J**). These results indicate that abolishing bacterial glycolysis leads to *Salmonella* dormancy in epithelial cells, which together with our previous results highlights how the interaction between host and bacterial metabolism are at play for successful infection.

## Discussion

During the infection of cells, *Salmonella* colonizes various intracellular niches and exhibits different growth rates (reviewed in ^25^). When infecting epithelial cells, it actively subverts host cells through the injection of bacterial effectors by T3SS-1^26^. *Salmonella* uptake can lead to the formation of the SCV, a unique endocytic compartment that is subsequently modified into a replicative niche through the action of effectors secreted by the T3SS-2. Inside the SCV *Salmonella* is able to grow, albeit the rate slows down overtime (**Figure S4**). In epithelial cells, a *Salmonella* subset ruptures the SCV and undergoes hyper-replication in the host cell cytosol within one hour after internalization^4,5^. A third dormant subpopulation was described in epithelial cells, characterized for a prolonged non-replication phase (**Figure S4**). Dormant *Salmonella* have been observed in macrophages, fibroblasts, and more recently in epithelial cells^6,17,27^ and recognized as an important pathogenic strategy for *Salmonella* as they persist after antibiotic treatment^28^.

To survive, intracellular bacteria need to adapt to the different environments and metabolism to replicate in their niche. Heterotroph organisms, such as bacteria, use organic carbon compounds as energy sources and substrates for all carbon intermediates. In epithelial cells and macrophages, *Salmonella* uses sugars, mainly glucose^21^, as the main carbon source (reviewed in^29^). Moreover, the role of the *Salmonella* glycolysis pathway for intracellular survival and replication in epithelial cells has been reported^21^. These results indicate that mutations in key glycolytic enzymes cause only a minor defect in intracellular replication, whereas glucose transport is essential for this process. Specifically, a triple mutant lacking all three glucose transport systems exhibits a significant impairment in intracellular replication^21^. Moreover, by using C^13^-isotopologue profiling, it was determined that *S*. Typhimurium uses glucose, but not glucose-6-P, as the preferential carbon substrate in Caco-2 cells. C^13^ was incorporated into amino acids, suggesting de novo synthesis from glucose by intracellular *Salmonella*^30^. All these studies are based on bulk assays, and as highlighted by our work, growth, and metabolism are tightly linked to the intracellular niche they occupy and need to be analyzed individually.

Some bacterial effectors have been associated with changes in intracellular niche formation. In particular, SifA and SopF have been associated with vacuolar escape and replication in the cytosol of epithelial cells^12,20,31^. It has been reported that in the absence of SifA, other effectors, namely SseJ and SopD2, destabilize the SCV membrane leading to increased cytosolic escape^16^. In the case of SopF, it has been determined that this effector contributes to vacuolar stability, by avoiding the recruitment of the xenophagy machinery via ADP-ribosylation of the vATPase^32^, with a significant increase in the proportion of Δ*sopF* bacteria present in the cytosol of HeLa cells^20^. In this work, we confirmed that τι*sifA* and τι*sopF* mutants have a higher chance of reaching the host cell cytosol, but this happens in only in less than 40% of infected cells (**Figure 1** and **S1**). This indicates that these effectors alter the balance between intracellular niches but they do not completely change the localization of invading *Salmonella*, as initially postulated for *sifA*^31^.

As expected from our previous data using a τι*ssaV* mutant^6^, deletion of T3SS-2 secreted effectors does not induce a significant change in terms of dormancy at 6 hpi. On the other hand, when we analyze a later time point we can observe an amplification of some of the cellular phenotypes that could be at play at early time points (**Figure 1** and **S1**), leading to widely different levels of formation of dormant *Salmonella*. We decided to focus on the mutant with the biggest increase in dormancy formation at this time point, τι*sifA*, as the effector SifA is implicated in the formation of the SIF network.

While the role of the SIF network is not completely elucidated, it has been hypothesized that allows *Salmonella* to gain access to endocytosed nutrients to promote intravacuolar proliferation, as labeled endosomal cargo can be detected within the SIF network of infected cells, demonstrating the access of these structures to endocytosed material^9,33^. However, it is very possible that the increase in the membrane surface available due to SIF presence also increases the recruitment of host nutrient transporters, contributing to *Salmonella* intravacuolar replication. Our data indicates that the metabolic state of the cell is highly relevant to avoid a dormant state, therefore it is plausible that a cell with a high glycolytic activity would lead to higher *Salmonella* replication due to scavenging of nutrients directly from the infected cell. In this work we have confirmed the contribution of SIFs to intravacuolar nutrition, but moreover, *Salmonella* enclosed in vacuolar compartments devoid of SIFs will end up in a metabolically stunted state, leading to dormancy.

Altogether our results indicate that the nutrient availability and the ability of *Salmonella* to properly hijack the host-cell endocytic pathway for intravacuolar nutrition are relevant during infection of epithelial cells. Moreover, the glycolytic state of the cell and *Salmonella*’s ability to utilize the glycolysis pathway are both key to avoiding the formation of dormant bacteria. This is particularly important in the context of infection and relapse of vulnerable and malnourished individuals, as it could lead to bacteria avoiding clearance by the immune system and the possibility of relapse.

## Acknowledgments

We thank Dr. Michael Hensel for kindly providing HeLa LAMP1-GFP cells. We are indebted to Dr. Jay Hinton and Dr. Roi Avraham for fruitful discussions about this work. We acknowledge the technical support in flow cytometry experiments from the Flow cytometry platform at Institut Pasteur, especially Sandrine Schmutz and Sebastien Meghabara. Work in the unit of J.E. is supported by the ERC (CoG Endosubvert) and the ANR (grants HBP sensing, PureMagRupture, and RabReprogram) and J.E. is a member of the LabEx “IBEID” and “Milieu Interieur”.T.M was supported by a Master’s students on AMR fellowship from Institut Pasteur. This research was supported by fellowships from Croucher Foundation (HK) and Fondation pour la Recherche Médicale (FR) to C.H.L. C.H.L. is part of the Pasteur - Paris University (PPU) International PhD Program.

## Author contributions

Conceptualization: CV and JE. Data curation: CV and FJGR. Formal analysis: CV, FJGR, TM, KD. Funding acquisition: JE. Investigation: CV, FJGR, TM, MG, KD. Methodology: CV and FJGR. Project administration: CV and JE. Resources: CV, FJGR, CHL. Software: FJGR. Supervision: CV and JE. Validation: CV and FJGR. Visualization: CV with help from FJGR. Writing – original draft: CV. Writing – review & editing: CV, FJGR and JE.

## Material and Methods

### Bacterial strains

All *Salmonella* strains used in this work are derived from the parental strain *S*. Typhimurium SL1344 and are listed in **Table S1**. Gene deletion mutants were obtained using the All mutants were constructed using the λ red recombinase system^34^ with modifications^35^ using the primers described in **Table S2**. For the τι*sopE*τι*sopE2* double mutant, the τι*sopE*::FRT deletion mutant was generated using plasmid pCP20^36^, and the elimination of the resistance cassette was verified by PCR. Then, the τι*sopE2*::Kan mutation was introduced by generalized transduction of P22 phage lysates^37^ and confirmed by PCR. All plasmids used in this study are described in **Table S3**. Bacteria were routinely cultured in Lysogeny broth (LB) supplemented with ampicillin 100 μg/mL or kanamycin 50 μg/mL, when needed.

### Mammalian cell culture

HeLa and HeLa LAMP1-GFP^7^ cells, a kind gift from Dr. Michael Hensel, were routinely cultured in Dulbecco’s Modified Eagle Medium (DMEM, high glucose, GlutaMAX, ThermoFisher) containing 10% (v/v) heat-inactivated fetal bovine serum (FBS, Sigma) and grown at 37 °C with 5% CO2 and 100% humidity. Cells were routinely tested for mycoplasma.

For all flow cytometry experiments, 3 X 10^5^ cells/well were seeded in 6-well plates 2 days prior to the infection in DMEM, high glucose, supplemented with GlutaMAX. For glycolysis inhibition experiments, the media was changed to DMEM no glucose supplemented with galactose 10mM and 10% FBS 24 h prior to infection, or to DMEM high glucose supplemented with 2DG 1mM and 10% FBS 3 h prior to infection.

### Salmonella infections

Bacteria strains were streaked from glycerol stock on LB agar plates with appropriate antibiotics 2 days prior to infection. Three bacterial colonies were used to inoculate an overnight (O/N) culture in LB medium supplemented with 0.3 M NaCl and the appropriate antibiotics and grown with shaking at 37 °C. The next day, 150 μL of this culture were sub-cultured in 3mL LB +0.3 M NaCl (1:20 dilution) and grown with shaking at 37 °C for 3h. Bacteria were harvested with centrifugation (1 mL, 6000 x g, 1 min, RT), washed once in 1x PBS and resuspended in DMEM without FBS (for high and low glucose comparisons, DMEM no glucose was used and supplemented with 25mM or 2.8mM glucose. For glycolysis inhibition experiments, DMEM no glucose supplemented with galactose 10mM or DMEM high glucose supplemented with 2-DG 1mM were used. HeLa cells were infected at an MOI of 100 bacteria/cell for 30 min at 37 °C. Extracellular bacteria were removed and washed 5 times with PBS. Infected cells were then incubated in DMEM +10% FBS for 1h, washed 3 times with PBS at 1 hpi and 3 hpi and then incubated in DMEM +10% FBS supplemented with 10 μg/mL gentamicin for the remaining time of the infection.

### Flow cytometry

At 6 and 24 hpi, cells were washed once with PBS and detached with 0.05% Trypsin for 5 min at 37 °C. Detached cells were mixed with equal volume of DMEM +10% FBS, filtered through a 40 μm strainer and collected by centrifugation (500 x g, 5 min, 4 °C). Cell pellets were dislodged and fixed in 4% PFA for 15 min at RT. Fixed cells were washed twice with PBS and finally resuspended in 500 μL PBS for further analysis. The fluorescence intensities of the samples were assayed with LSR Fortessa (BD) (tagBFP Ex: 405 nm Em: 450/50 nm; Timer510 Ex: 488 nm Em: 525/50 nm; Timer580 Ex: 562 nm Em: 582/15 nm; smURFP Ex: 633 nm Em: 670/30 nm) and analyzed with FlowJo (v10.10.0). For each sample, a minimum of 10,000 cells were analyzed. The recorded events were gated according to the strategy described in **Figure S1-A**.

### Live confocal microscopy for SIF analysis

For live-imaging confocal microscopy, HeLa LAMP1-GFP cells^7^ were seeded in 8-well ibidi μslides (80827-90) at a density of 10,000 cells/well 2 days prior to infection. Cells were infected as described above. When needed, 1 μM nocodazole (M1404, Sigma), or the equivalent volume of DMSO, was added to the infected cells at 1 hpi, and maintained throughout the experiment. At 6 hpi, cells were washed, and media was replaced with FluoroBrite DMEM (A1896701, Gibco) supplemented with 10% FBS, 10 mM HEPES (15630-056, Gibco), GlutaMAX (35050-038, Gibco), Sodium pyruvate (11360-039, Gibco) and 10 μg/mL Gentamicin (G1272, Sigma).

### Staining and automated time-lapse confocal imaging acquisition

For automated time-lapse confocal imaging experiments. We performed infection assays in 384-well microplates. HeLa and LAMP1-GFP HeLa cells were seeded 24 hours before infection at a density of 8,000 cells per well. Each experiment was performed with four technical replicates and repeated to obtain at least three biological replicas. To track individual cells, nuclei and cytoplasm were stained one hour before infection with 200 ng/mL Hoechst H33342 (H3570, Life Technologies) and 25 µM CellTracker Blue (C12881, Invitrogen), respectively. Cells were washed before infection to prevent bacterial staining. Infections with *S*. Typhimurium strains carrying SINA 1.1 or SINA 1.7, as well as treatments with nocodazole, were conducted as previously described. Image acquisition was performed using an automated microlens-enhanced spinning disc confocal microscope (Opera Phenix High Content Screening System, PerkinElmer) equipped with a 63x water objective. Excitation lasers operated at 405, 488, 561, and 640 nm, with emission filters set at 450, 540, 600, and 690 nm. To capture the dynamics of infection, images of multiple fields (ranging from 10 to 16) were acquired every 30 minutes over a 9-hour period in an incubation chamber maintained at 37°C with 5% CO2.

### Image analysis of automated time-lapse confocal imaging

Single-cell and single-bacteria analyses were conducted using Harmony software v.4.9 (PerkinElmer) and custom-built scripts. For HeLa cells infected with *S*. Typhimurium carrying SINA 1.1, we segmented cell nuclei using the Hoechst signal and cytoplasm using the CellTracker Blue background. Hoechst provided a strong nuclear signal, while CellTracker Blue generated a faint cytoplasmic signal, enabling the detection of both nuclei and cytoplasm within a single microscope channel (405/450). Intracellular bacteria were segmented by measuring the Timer510 and Timer580 signal intensities in the 488/540 and 561/600 channels, respectively. Each bacterium was then subclassified as “dormant,” “cytosolic,” or “vacuolar” based on their Timer510/580 ratio, smURFP, and tagBFP intensity, respectively. For LAMP1- GFP HeLa cells infected with *S*. Typhimurium carrying SINA 1.7, cytoplasm segmentation was achieved by combining the intensity signals of LAMP1-GFP and CellTracker Blue. Intracellular bacteria were segmented based on the dsRED reporter signal intensity in the 561/600 channel. To quantify SIFs at the single-cell level, we utilized an interactive training mode (PhenoLOGIC™) integrated into Harmony software v.4.9. This approach enabled the identification of regions within the LAMP1-GFP signal based on variations in texture. Specifically, we defined a texture region that accurately corresponded to SIFs while distinguishing it from regions with globular distributions or lacking LAMP1. To minimize false positives, we conducted morphological analyses of SIFs and established cut-off criteria based on their size and length. Cells containing at least one SIF were classified as positive, and this measurement was applied to all infected cells

### Immunofluorescence microscopy

Cells seeded on coverslips were infected as described above. At selected time points, cells were washed with PBS and fixed in 4% PFA for 10 min at RT. After washing 3 times with PBS, free aldehyde groups were quenched using 50mM NH_4_Cl in PBS for 10 min at RT. Permeabilization and blocking were performed using 0.1 % Saponin, 5 % donkey normal serum, and 1% BSA in PBS for 30 min at RT. Coverslips were washed 3 times with PBS and incubated with rabbit recombinant monoclonal anti-RAB7 (Abcam # ab137029) in 1:100 dilution in blocking buffer O/N at 4 °C. The next morning, coverslips were washed 3 times with PBS and incubated with Alexa Fluor-488 polyclonal donkey anti-rabbit IgG (H+L) (catalog # A-21206) at 1:400 dilution in blocking buffer for 1h at RT. Finally, coverslips were mounted on SuperFrost Plus microscope slides (Thermo Scientific) with Fluoromount-G Mounting Medium (Invitrogen). Samples were imaged on a NIKON Eclipse Ti2-E Spinning Disk Confocal Microscope, equipped with a Yokogawa CSU-W1 Confocal Scanner Unit and an NSPARC Detector unit. Images were analyzed with FIJI (NIH) and Amira software (ThermoScientific), and figures were prepared using Inkscape (v1.0.2) and Biorender.

## Supplementary figures

**Figure S1.**
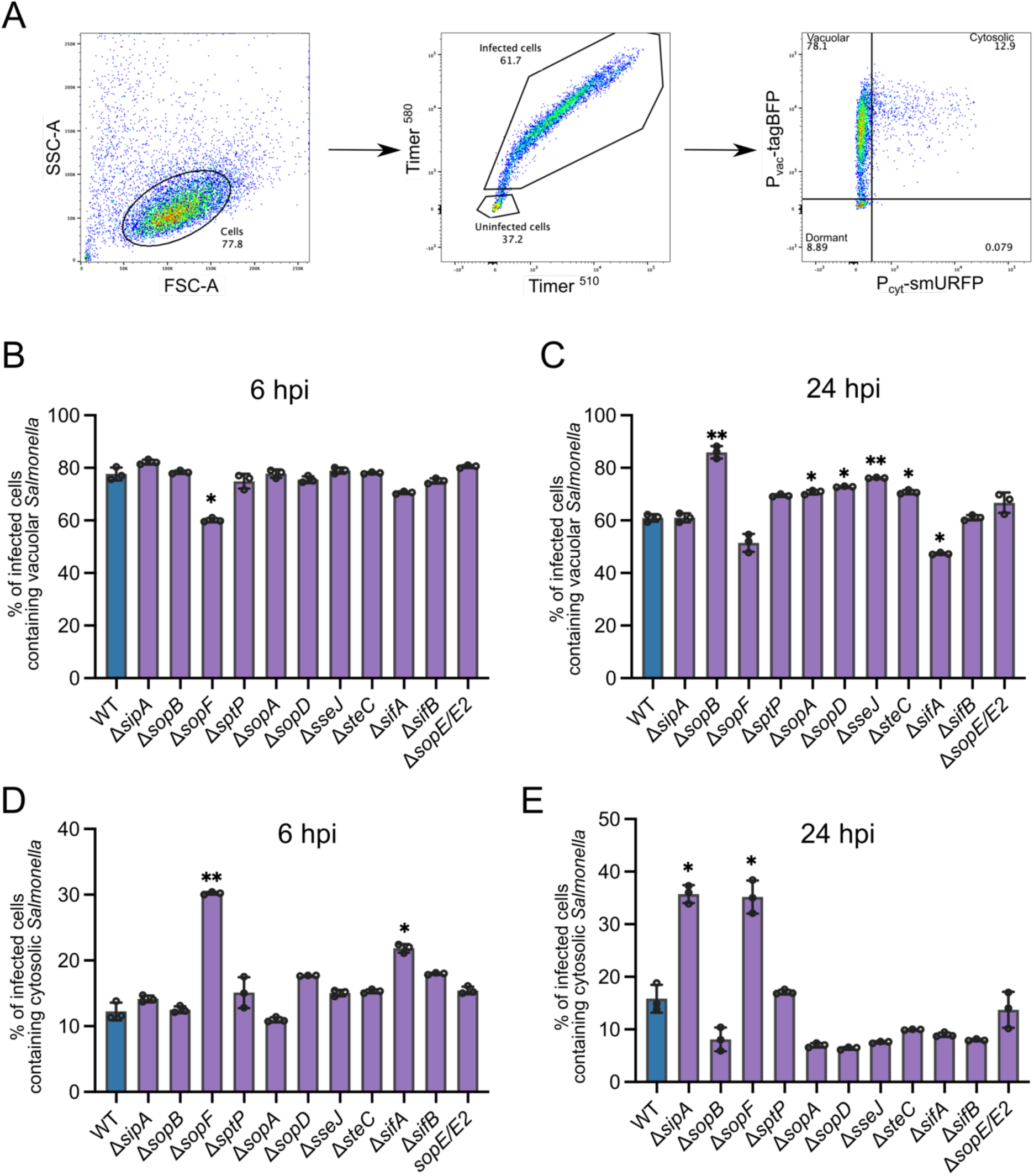
*Salmonella* effector mutants show altered intracellular niche formation, related to Figure 1. **(A)** Gating strategy used for all analysis of epithelial cells infected with *Salmonella* expressing SINA1.1. The percentage of cells containing vacuolar *Salmonella* at 6 **(B)** and 24 hpi **(C)** or cytosolic *Salmonella* at 6 **(D)** and 24 hpi **(E)** of the different deletion mutants (purple) compared with the WT strain (blue) is shown. Plots show the average of 3 biological replicates (N=3). Statistical significance was evaluated using a one-way ANOVA followed by a with Dunnett’s multiple comparison test, * = *P*<0.05, ** = *P*<0.01.

**Figure S2.**
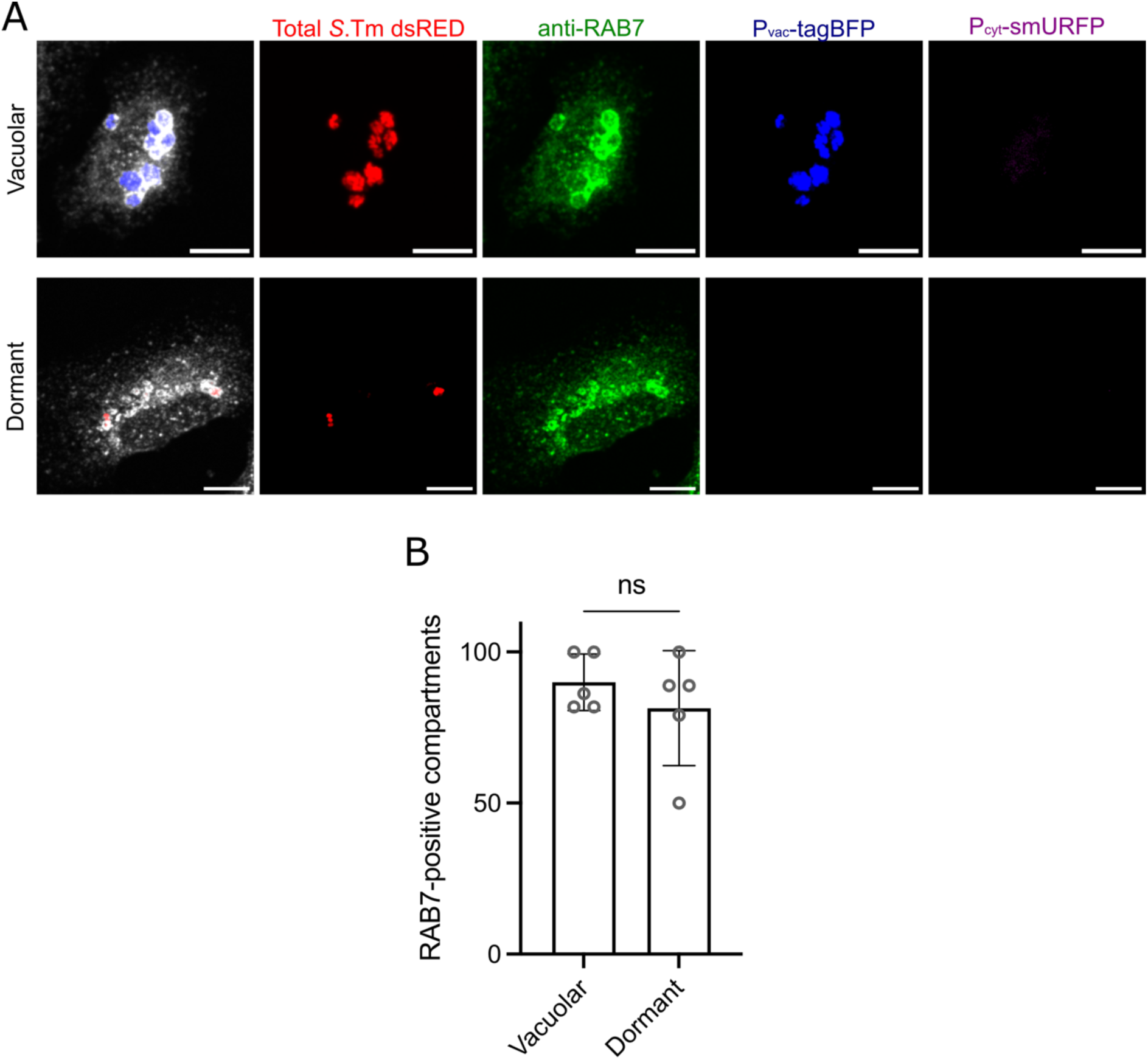
Dormant *Salmonella* is contained in a RAB7-positive compartment. **(A)** Representative confocal images of HeLa cells infected with *Salmonella* expressing SINA1.7 growing in a SCV (upper panel) or in a dormant state (lower panel). Green: immunofluorescence staining using a rabbit anti-RAB7 antibody followed by Alexa Fluor-488 anti-rabbit antibody, red: dsRED total *Salmonella*, blue: vacuolar marker P_vac_-tagBFP, magenta: cytosolic marker P_cyt_-smURFP. Merge shows the RAB7 distribution and *Salmonella* according to its intracellular niche (blue: vacuolar, red: dormant). **(B)** Quantification of the percentage of RAB7-positive compartments from 4 biological replicates (N=4). A total of 115 cells were analyzed with a total number of bacterial clusters for vacuolar (n=354) and dormant (n=152). Statistical significance was evaluated using an unpaired *t*-test with Welch’s correction.

**Figure S3.**
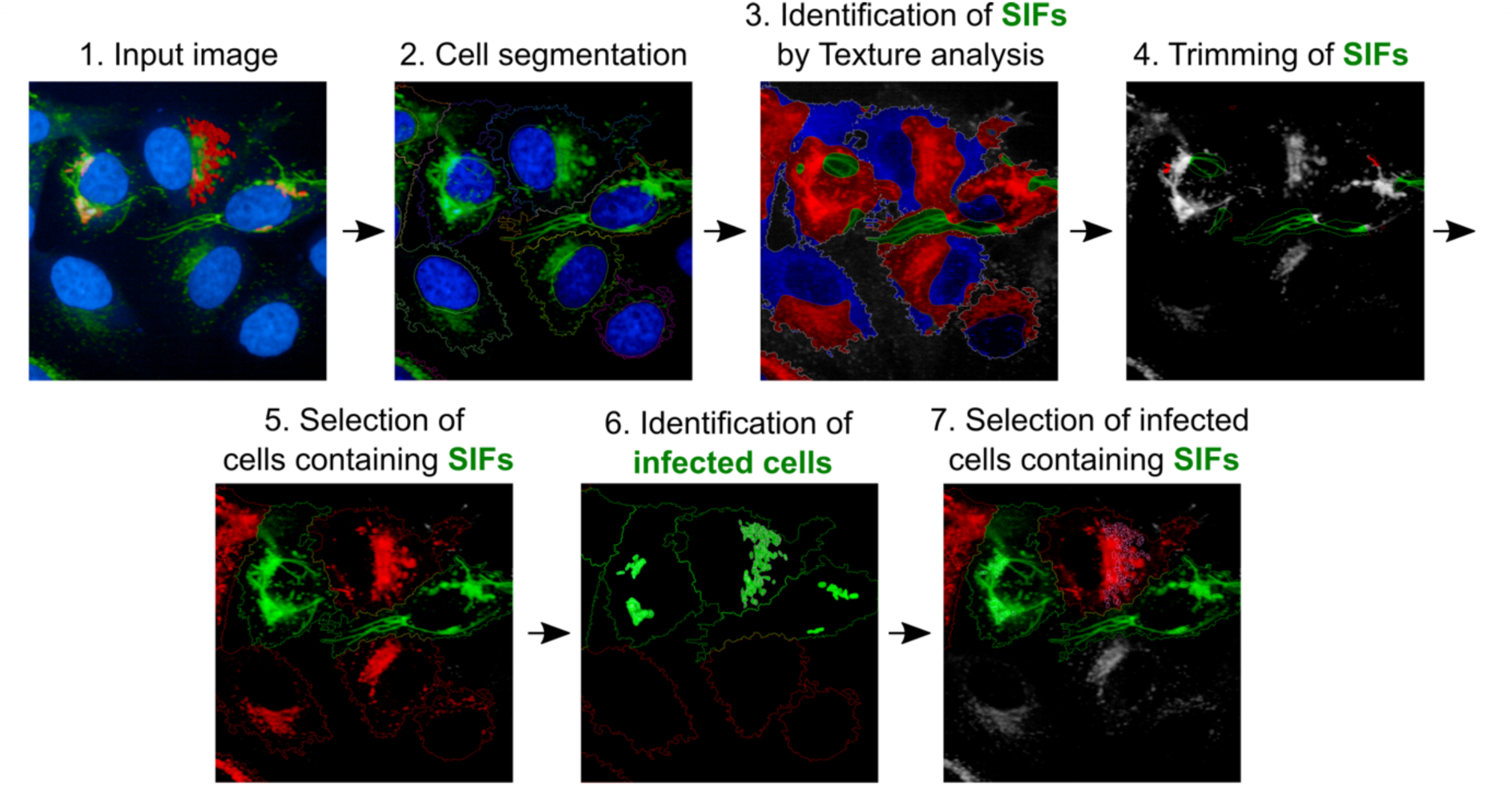
Schematic representation of the automated image analysis to track SIF formation in HeLa cells infected with *Salmonella*. HeLa cells expressing LAMP1-GFP were infected with *Salmonella* expressing SINA 1.7 (red), stained with Hoechst (blue), and visualized using confocal microscopy. High-content analysis (HCA) was performed on high-resolution raw images, which were filtered and automatically segmented to detect nuclei and cytoplasm. Texture analysis of the green channel was conducted using the interactive training mode of PhenoLOGIC™ to identify LAMP1 filamentous structures (SIFs). To minimize false positives, small LAMP1 filaments were excluded. Subsequently, cells were filtered based on the presence of SIFs and intracellular bacteria. Scale bar: 20 μm.

**Figure S4.**
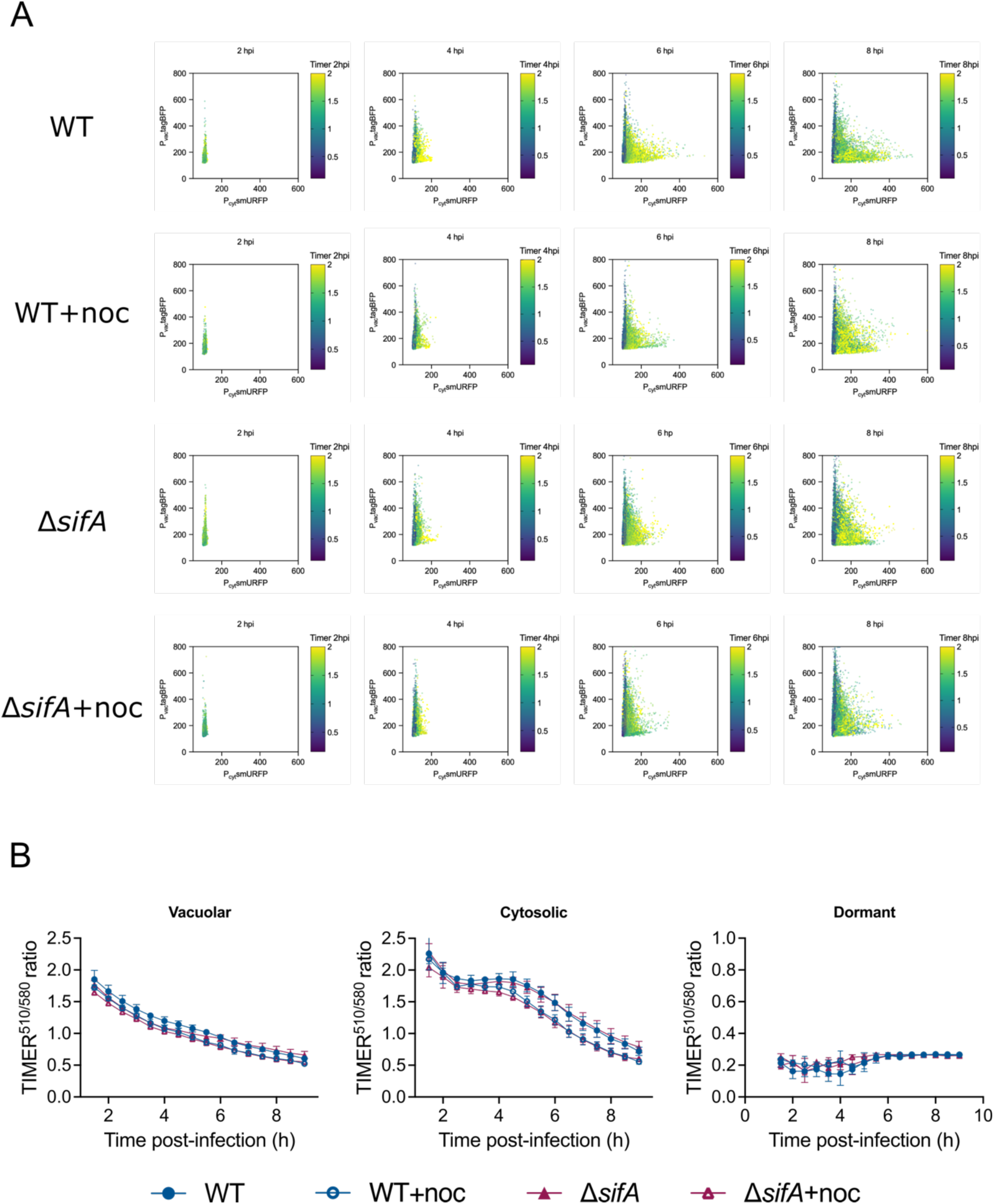
Automated image-based analysis of intracellular niche formation: **(A)** Single-bacteria tracking of the presence of vacuolar (P_vac_-tagBFP^+^), cytosolic (P_cyt_-smURFP^+^) and dormant (P_vac_-tagBFP^-^/ P_cyt_-smURFP^-^) at different time points, and their respective Timer 510/580 ratio over time. **(B)** Intracellular growth rate as a function of the Timer 510/580 ratio for each subpopulation over time in treated and untreated cells.

