## Supplementary material for "SifA-mediated Remodeling of the *Salmonella*-Containing Vacuole Prevents Bacterial Dormancy by Promoting Nutrient Accessibility": Table S1

**Table S1**: Strains used in this study

| Strain | Genotype |
|---|---|
| Δ*sipA* | Δ*sipA*::Kan |
| Δ*sopB* | Δ*sopB*::Kan |
| Δ*sopF* | Δ*sopF*::Kan |
| Δ*sptP* | Δ*sptP*::Kan |
| Δ*sopA* | Δ*sopA*::Kan |
| Δ*sopD* | Δ*sopD*::Kan |
| Δ*sseJ* | Δ*sseJ*::Kan |
| Δ*steC* | Δ*steC*::Kan |
| Δ*sifA* | Δ*sifA*::Kan |
| Δ*sifB* | Δ*sifB*::Kan |
| Δ*sopE/E2* | Δ*sopE::FRT* Δ*sopE2::Kan* |
| Δ*pfkA* | Δ*pfkA*::Kan |
| Δ*pfkB* | Δ*pfkB*::Kan |
| Δ*pfkAB* | Δ*pfkA::FRT* Δ*pfkB::Kan* |
| Δ*manZ* | Δ*manZ*::FRT |
| Δ*manZ* Δ*sifA* | Δ*manZ::FRT* Δ*sifA::Kan* |
