## Supplementary material for "SifA-mediated Remodeling of the *Salmonella*-Containing Vacuole Prevents Bacterial Dormancy by Promoting Nutrient Accessibility": Table S2

**Table S2**: Primers used in this study

| Name | Sequence |
|---|---|
| sipA_H1+P1 | GATATTAATAATGGTTACAAGTGTAAGGACTCAGCCCCCCGTGCAGGCTGGAGCTGCTTC |
| sipA_H2+P2 | TCCCGGTTAATTAACGCTGCATGTGCAAGCCATCAACGGTCATATGAATATCCTCCTTAG |
| sipA_Out5 | CAGACGCTGACGCAAAAATA |
| sipA_Out3 | GATCCTCAACCAGATGGGTC |
| sopB_H1+P1 | TAAAAACGCTATGCAAATACAGAGCTTCTATCACTCAGCTGTGCAGGCTGGAGCTGCTTC |
| sopB_H2+P2 | ACCTCAAGACTCAAGATGTGATTAATGAAGAAATGCCTTTCATATGAATATCCTCCTTAG |
| sopB_Out5 | AGAGACAAAAGCGGCAAAAA |
| sopB_Out3 | GGCATAAAGGGACAGCACAT |
| sopF_H1+P1 | CAGGAGACATATGCTCAAACCTATCTGCCATAGTGGAAGTGTGCAGGCTGGAGCTGCTTC |
| sopF_H2+P2 | AATAAGCTTGTCAATATAATATTATGCAGTCTCTATTAAGCATATGAATATCCTCCTTAG |
| sopF_Out5 | TGAGTTTGGTGACGCGATTA |
| sopF_Out3 | ACAGGTATCTGCCAGAACGG |
| sptP_H1+P1 | CTGCAGGAATATGCTAAAGTATGAGGAGAGAAAATTGAATGTGCAGGCTGGAGCTGCTTC |
| sptP_H2+P2 | TATGTTTTTATCAGCTTGCCGTCGTCATAAGCAACTGGGCCATATGAATATCCTCCTTAG |
| sptP_Out5 | TGATATGTGTTCCGATGCGT |
| sptP_Out3 | GGAATGTCAGCAGAAGAGAAAAA |
| sopA_H1+P1 | AGGAATTCTAATGAAGATATCATCAGGCGCAATTAATTTTGTGCAGGCTGGAGCTGCTTC |
| sopA_H2+P2 | TGAGGCTGGACTACGCCCAGGCCAGTGGCAGGATGGATGACATATGAATATCCTCCTTAG |
| sopA_Out5 | ACGGTGACATGATGACAGGA |
| sopA_Out3 | TATGGAATTTAGGCCAGGGG |
| sopD_H1+P1 | GGAAAATATTATGCCAGTCACTTTAAGCTTCGGTAATCATGTGCAGGCTGGAGCTGCTTC |
| sopD_H2+P2 | CTGACTATCTTTATGTCAGTAATATATTACGACTGCACCCCATATGAATATCCTCCTTAG |
| sopD_Out5 | CGTCGAGACTTTCCCCAATA |
| sopD_Out3 | GAAGAGACACGCTTCTTCGG |

| | |
|---|---|
| sopE_H1+P1 | GATCATTACCGTGACAAAAATAACTTTATCTCCCCAGAATGTGCAGGCTGGAGCTGCTT |
| sopE_H2+P2 | TTTTCAGTGTTCAGGGAGTGTTTTGTATATATTTATTAGCCATATGAATATCCTCCTTAG |
| sopE_Out5 | CAAGCAACCGTCCGGCCTGC |
| sopE_Out3 | AAAAGCGGAACTTCTTGCTG |
| sopE2_H1+P1 | GAGAACTACCGTGACTAACATAACACTATCCACCCAGCACGTGCAGGCTGGAGCTGCTTC |
| sopE2_H2+P2 | TTTACTACCATCAGGAGGCATTCTGAAGATACTTATTCGCCATATGAATATCCTCCTTAG |
| sopE2_Out5 | CTGTTCAGGATTGTCCCGAT |
| sopE2_Out3 | GTTTTGTTAAAAGCGTCGCC |
| sseJ_H1+P1 | GGAGGACACTATGCCATTGAGTGTTGGACAGGGTTATTTCGTGCAGGCTGGAGCTGCTTC |
| sseJ_H2+P2 | CGATGGAACTTTATTCAGTGGAATAATGATGAGCTATAAACATATGAATATCCTCCTTAG |
| sseJ_Out5 | CTCACGCCAGCACACTAAAA |
| sseJ_Out3 | ATCGGCAGCAAAGATAGCAT |
| steC_H1+P1 | GATGAGACATATGCCGTTTACATTTCAGATCGGAAATCATGTGCAGGCTGGAGCTGCTTC |
| steC_H2+P2 | GAACTAAATGCTATTTTTTTAATTCATCCTTTAATACCTTCATATGAATATCCTCCTTAG |
| steC_Out5 | CACACGGTAACGAAGTTCCT |
| steC_Out3 | GCTACAGGCTGTCCAGATCC |
| sifA_H1+P1 | TGAGATTAATATGCCGATTACTATAGGGAATGGTTTTTTAGTGCAGGCTGGAGCTGCTTC |
| sifA_H2+P2 | TCGTCTGATTTTATAAAAAACAACATAAACAGCCGCTTTGCATATGAATATCCTCCTTAG |
| sifA_Out5 | ATCCGCGGTAGTCCTTCTTT |
| sifA_Out3 | TATTGTGCCTGGCAAGAGGT |
| sifB_H1+P1 | GGTCTACATTATGCCAATTACTATCGGGAGAGGATTTTTAGTGCAGGCTGGAGCTGCTTC |
| sifB_H2+P2 | TATGGTGTGATCAACTCTGGTGATGAGCCTCATTTTTTGTCATATGAATATCCTCCTTAG |
| sifB_Out5 | CTTCATTTTGAGCCTCCTCG |
| sifB_Out3 | ACGAGCCAATTCGTTCCATA |
| pfkA_H1+P1 | AGAGGTAGTCATGATTAAGAAAATCGGTGTGTTGACAAGCGTGCAGGCTGGAGCTGCTTC |
| pfkA_H2+P2 | GCAAAAACAATCAGTACAGTTTTTTCGCGCACTCCATCCACATATGAATATCCTCCTTAG |
| pfkA_Out5 | GGGGTTATCCTGGTACGGTT |

| | |
|---|---|
| pfkA_Out3 | ATAAATCTGAGGGATTGCCG |
| pfkB_H1+P1 | GGAGGTAACGATGGTACGTATCTATACGTTGACGCTTGCGGTGCAGGCTGGAGCTGCTTC |
| pfkB_H2+P2 | GGGGAAACGATTATTGCGCGGAAAGATAGGCGTATATTTTCATATGAATATCCTCCTTAG |
| pfkB_Out5 | CTGACTGAGCAAGGTAGCCC |
| pfkB_Out3 | AACGGGTAATATCGCCACTG |
| manZ_H1+P1 | GGTGAGCGAAATGGTTGATATGACTAAAACTACCACCGAGGTGCAGGCTGGAGCTGCTTC |
| manZ_H2+P2 | TACAACAGCCTTACTGGCCCAGCAGTCCTACGGAGTAGCCCATATGAATATCCTCCTTAG |
| manZ_Out5 | CAGGCCAGGTACTGACCATT |
| manZ_Out3 | TGCGGCAATAAAGAGAATCA |
