## Supplementary material for "SifA-mediated Remodeling of the *Salmonella*-Containing Vacuole Prevents Bacterial Dormancy by Promoting Nutrient Accessibility": Table S3

**Table S3**: Plasmids used in this study

| Name | Characteristics or GenBank accession number | Reference |
|---|---|---|
| SINA1.1 | pBR322::$P_{const}$-TIMER BAC, $P_{vac}$-tagBFP and $P_{cyt}$-smURFP; $Amp^R$ | 6 |
| SINA1.7 | pBR322::$P_{const}$-dsRED, $P_{vac}$-tagBFP and $P_{cyt}$-smURFP; $Amp^R$ | 6 |
| pKD46 | AY048746.1 | 34 |
| pCP20 | FLP+, ƛ cI857+, ƛ$P_R$ $Rep^{ts}$, $Amp^R$, $Cam^R$ | 36 |
| pCLF4 | HM047089 | 35 |
